# Orthogonal CRISPR-Cas tools for genome editing, inhibition, and CRISPR recording in zebrafish embryos

**DOI:** 10.1101/2020.11.07.372151

**Authors:** Paige R. Takasugi, Shengzhou Wang, Kimberly T. Truong, Evan P. Drage, Sahar N. Kanishka, Marissa A. Higbee, Nathan Bamidele, Ogooluwa Ojelabi, Erik J. Sontheimer, James A. Gagnon

## Abstract

The CRISPR-Cas universe continues to expand. The type II CRISPR-Cas system from *Streptococcus pyogenes* (SpyCas9) is the most widely used for genome editing due to its high efficiency in cells and organisms. However, concentrating on a single CRISPR-Cas system imposes limits on target selection and multiplexed genome engineering. We hypothesized that CRISPR-Cas systems originating from different bacterial species could operate simultaneously and independently due to their distinct single-guide RNAs (sgRNAs) or CRISPR-RNAs (crRNAs), and protospacer adjacent motifs (PAMs). Additionally, we hypothesized that CRISPR-Cas activity in zebrafish could be regulated through the expression of inhibitory anti-CRISPR (Acr) proteins. Here, we use a simple mutagenesis approach to demonstrate that CRISPR-Cas systems from *Streptococcus pyogenes* (SpyCas9), *Streptococcus aureus* (SauCas9), *Lachnospiraceae bacterium* (LbaCas12a, previously known as LbCpf1), are orthogonal systems capable of operating simultaneously in zebrafish. CRISPR systems from *Acidaminococcus* sp. (AspCas12a, previously known as AsCpf1) and *Neisseria meningitidis* (Nme2Cas9) were also active in embryos. We implemented multichannel CRISPR recording using three CRISPR systems and show that LbaCas12a may provide superior information density compared to previous methods. We also demonstrate that type II Acrs (anti-CRISPRs) are effective inhibitors of SpyCas9 in zebrafish. Our results indicate that at least five CRISPR-Cas systems and two anti-CRISPR proteins are functional in zebrafish embryos. These orthogonal CRISPR-Cas systems and Acr proteins will enable combinatorial and intersectional strategies for spatiotemporal control of genome editing and genetic recording in animals.

## Introduction

The use of clustered regularly interspaced short palindromic repeats (CRISPR) and CRISPR associated proteins (Cas) for genome editing has expanded significantly in recent years. CRISPR-Cas systems have several advantages over previous systems, such as zinc-finger nucleases and transcription activator-like effector nucleases, including the ease of design and use, low cost, high efficiency, and customizability (Adli 2018; Knott and Doudna 2018; K. Liu et al. 2019). A large variety of CRISPR-Cas systems have been described, originating from different bacterial species. Currently, these systems are organized into two large classes and further divided into six types based on the unique *cas* genes they contain (Makarova et al. 2019). Class 1 systems use multiple Cas proteins in the effector complex, while Class 2 utilize a single protein for endonuclease activity. Class 2 systems are most commonly used for genome editing, usually type II and type V, due to the ease of delivering a single multi-domain protein in eukaryotes. Type II and type V systems have a few notable differences, including the enzyme and guide RNA structures, their DNA target sequences, and the manner in which they create double-stranded breaks (**Figure 1A, B**). These characteristics can be significant for genome editing if they result in different editing outcomes or permit targeting to different regions of the genome.

**Figure 1:**
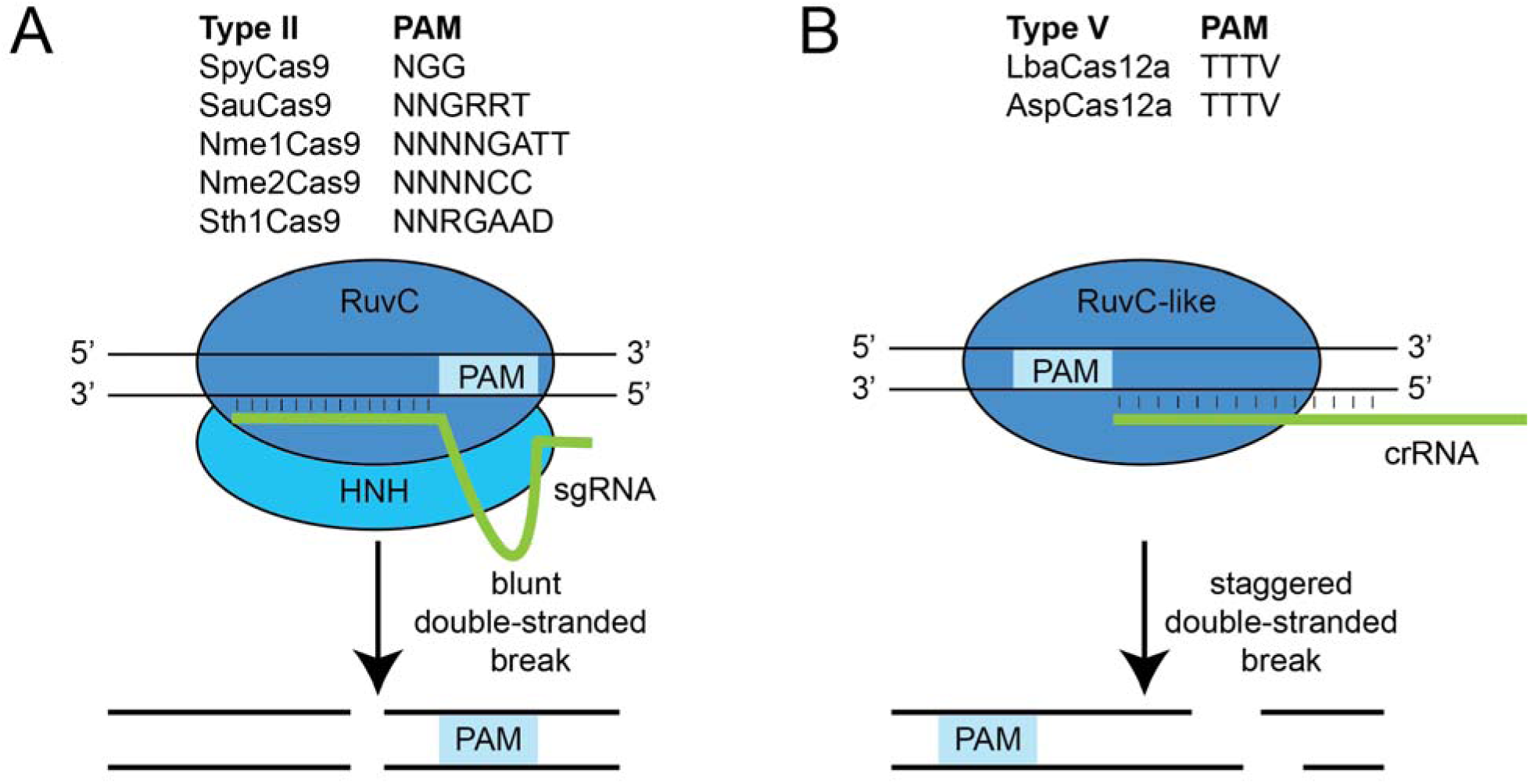
An overview of Type II and V CRISPR-Cas systems. **A**. Type II CRISPR-Cas systems employ a multidomain protein (Cas9) which complexes with CRISPR RNA (crRNA) and trans-activating CRISPR RNA (tracrRNA) to cause target DNA cleavage. These two RNA molecules can be fused into a single-guide RNA (sgRNA), as shown. The Cas9 HNH domain cleaves the complementary strand, while the RuvC domain cleaves the non-complementary strand in the same position. This results in a blunt double-strand break. **B**. Type V CRISPR-Cas systems employ a distinct multidomain protein (Cas12a, previously known as Cpf1), which complexes with and processes a crRNA to target DNA for cleavage. Cas12a does not require a tracrRNA. Type V enzymes contain a RuvC-like domain, but do not have an HNH nuclease domain (Zetsche et al. 2015; Makarova et al. 2019). This RuvC-like domain cleaves both DNA strands to create a staggered double-strand break. In either case, double-strand breaks can be repaired by the cells using a variety of DNA repair mechanisms, often resulting in insertions or deletions (indels). PAMs for each system are shown above the diagram.

The zebrafish has historically been a testbed for reverse genetic and RNA knockdown tools in animals, mainly due to regular access to large numbers of fertilized eggs that are easy to microinject (Nasevicius and Ekker 2000; Meng et al. 2008; Doyon et al. 2008; Huang et al. 2011; Bedell et al. 2012; Sander et al. 2011; Dahlem et al. 2012; Hwang et al. 2013; Jao, Wente, and Chen 2013; Gagnon et al. 2014; Feng et al. 2016; Moreno-Mateos et al. 2017; K. Liu et al. 2019).. Indeed, the widely-used SpyCas9 system was first demonstrated in zebrafish embryos before applications to other organisms (Hwang et al. 2013; Chang et al. 2013). Since its introduction, notable improvements to SpyCas9 genome editing in zebrafish include computational prediction of active sgRNAs, methods for rapid sgRNA generation, multiplexed editing, and the use of concentrated SpyCas9 protein and commercially-available crRNAs/tracrRNA (Gagnon et al. 2014; Moreno-Mateos et al. 2015; Shah et al. 2015; Varshney et al. 2015; Labun et al. 2016; Burger et al. 2016; Thyme et al. 2016; Wu et al. 2018; DiNapoli et al. 2018; Ata et al. 2018; Hoshijima et al. 2019; Kroll et al. 2020).

However, other CRISPR-Cas systems have been relatively underexplored in most model organisms, including zebrafish. While SpyCas9 has been widely used, a limited number of publications have described the activity of CRISPR systems from *Streptococcus aureus* (SauCas9), *Lachnospiraceae bacterium* (LbaCas12a), *Francisella novicida* (FnoCas12a), and *Acidaminococcus sp*. (AspCas12a) in zebrafish embryos (Feng et al. 2016; Moreno-Mateos et al. 2017; P. Liu et al. 2019). These systems expand the targetable space of the genome due to their distinct PAMs and may empower intersectional strategies that employ multiple CRISPR-Cas systems. Additionally, CRISPR-Cas type II systems create blunt double-stranded breaks while type V systems generate a staggered cut, which has implications for both indel and knock-in mutagenesis in zebrafish (Moreno-Mateos et al. 2017).

CRISPR systems have been adapted for biological recording, including in animals for lineage tracing (Farzadfard and Lu 2018; McKenna and Gagnon 2019; Wagner and Klein 2020). In these applications, expression of CRISPR components generates permanent “edits” to a barcode, recording lineage relationships or cell signaling pathway activation. These barcodes can be recovered using DNA or RNA sequencing. The technology is often limited by the ability to induce recording at just one or two time points with some crosstalk (Raj et al. 2018). Currently, nearly all applications have relied solely on SpyCas9 and have not explored alternative CRISPR systems. Expanding to additional CRISPR systems could enable new modes of temporal control over CRISPR recording, or multichannel recording of different features of cellular history.

Spatial and temporal control over genome editing in animals permit tissue-specific and developmental-stage specific mutagenesis for more sophisticated screens (Ablain et al. 2015; Yin et al. 2015; Shiraki and Kawakami 2018). These strategies generally rely on regulation of Cas enzyme expression, with ubiquitously-expressed guide RNA(s). However, these strategies can be leaky, have limited temporal control, and may require extensive molecular cloning and transgenesis. This limits their widespread adoption for medium- or large-scale genetic screens. A promising alternative strategy would employ anti-CRISPR (Acr) proteins (Marino et al. 2020). These proteins are capable of blocking CRISPR-Cas activity through direct interaction with Cas proteins, preventing DNA target site recognition or preventing DNA cleavage. Tissue- or time-specific expression of Acr proteins could provide an alternative approach to control over CRISPR-Cas genome editing (Lee et al. 2019). Although many Acr proteins have been identified and tested in bacteria and mammalian cell lines (Pawluk et al. 2016; Rauch et al. 2017), none have been validated in zebrafish.

Here, we implemented a simple assay to functionally assess CRISPR-Cas systems and inhibitory Acr proteins in zebrafish embryos. We found that CRISPR-Cas systems from SpyCas9, SauCas9 and LbaCas12a were highly active for F0 mutagenesis and are functionally orthogonal. We further established that orthologous CRISPR systems can be used for simultaneous genome editing in the same embryo. We also found that Nme2Cas9 was active in zebrafish embryos, albeit with limited efficacy relative to these other systems. We demonstrate that orthogonal CRISPR systems can be used for information-dense, multichannel CRISPR recording. Finally, we determined that Acr proteins can be effective inhibitors of SpyCas9 and SauCas9 in zebrafish. Together, these tools will enable sophisticated genome editing and CRISPR recording strategies in a variety of organisms.

## Results

### A simple assay for efficient CRISPR-Cas mutagenesis in zebrafish embryos

Many common quantitative assays exist to measure mutagenesis, such as the T7 Endonuclease 1 assay, RFLP mapping, Sanger sequencing or Illumina sequencing. These assays have many advantages, but can be expensive, require specialized equipment, and/or require a significant amount of molecular work. We implemented a simple phenotypic visual readout for CRISPR-Cas mutagenesis to allow screening of new candidate CRISPR systems. We designed single-guide RNAs (sgRNAs) for Cas9 systems, or CRISPR RNAs (crRNAs) for Cas12a systems, targeting multiple sites of the gene *tyrosinase (tyr). tyr* encodes an enzyme responsible for the conversion of tyrosine to melanin (Haffter et al. 1996; Camp and Lardelli 2001). Homozygous *tyr* mutant zebrafish embryos lack pigmentation, an easily observed phenotype at 2 or 3 dpf (days post-fertilization). For efficient F0 mutagenesis, we pooled together 3-5 sgRNAs or crRNAs, each targeting different sites in the *tyr* gene. A solution of CRISPR-Cas mRNA or protein and a pool of sgRNAs or crRNAs were microinjected into single-cell zebrafish embryos (**Figure 2A**). At 2 dpf, healthy embryos were screened for pigmentation loss and classified into one of four different categories: Not pigmented, mostly not pigmented, mostly pigmented and fully pigmented (defined as 0-5% pigmentation loss, 6-50% pigmentation loss, 51-99% pigmentation loss and 100% pigmentation loss, respectively) (**Figure 2B**). While this is not as sensitive as alternative assays, as it requires homozygous inactivation of *tyr*, our strategy is a rapid, easy, and cost-effective test for CRISPR-Cas functionality that requires no specialized equipment.

**Figure 2:**
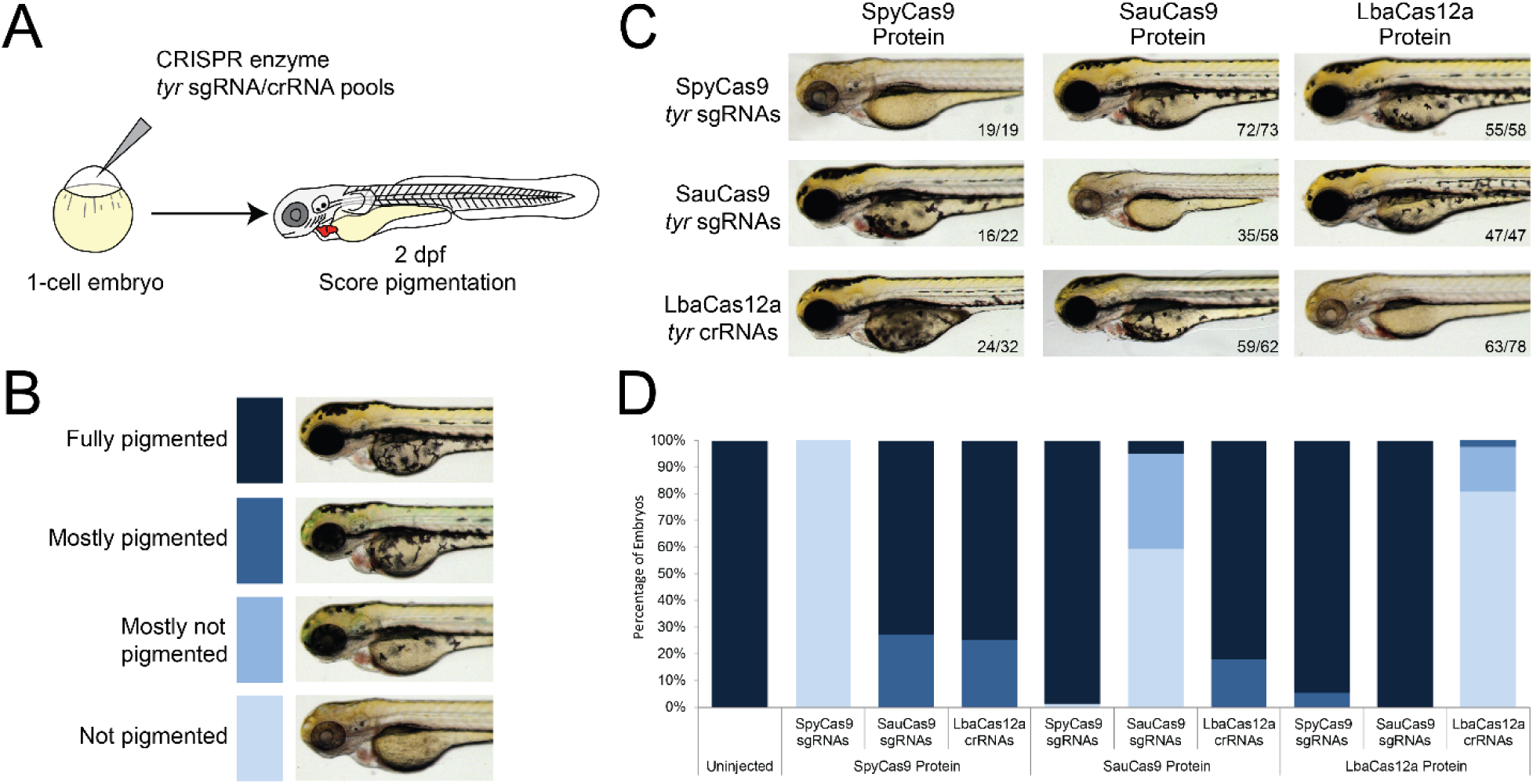
Highly efficient, orthogonal SpyCas9, SauCas9 and LbaCas12a genome editing in zebrafish embryos. **A**. Experimental design of the CRISPR screening method. A mix of CRISPR-Cas enzyme and *tyr* sgRNAs/crRNA pools is microinjected into the single-cell zebrafish embryo. Injected embryos are screened at 2 dpf for their level of pigmentation, an effective proxy for *tyr* mutagenesis. **B**. Example images of the four categories used to score pigmentation in embryos. Each category is roughly defined within a certain percentage of pigmentation: fully pigmented=100% pigmented, mostly pigmented=51-99% pigmented, mostly not pigmented=6-50% pigmented, not pigmented=0-5% pigmented. The associated colors act as the legend for panel D. **C**. Representative images of the phenotype of embryos after targeting the *tyr* gene with combinations of each CRISPR enzyme and pools of sgRNAs/crRNAs. Each CRISPR-Cas system is only functional when used with its corresponding sgRNAs/crRNAs. **D**. Quantification of pigmentation categories after microinjection as described in panel C. Raw data in Table S3. Panel B serves as the legend.

### Orthogonal and efficient CRISPR-Cas systems in zebrafish embryos

To screen for alternative CRISPR-Cas systems in zebrafish, we selected a variety of systems with unique protospacer adjacent motifs (PAMs). These included CRISPR-Cas systems from *Streptococcus pyogenes* (SpyCas9), *Streptococcus aureus* (SauCas9), *Streptococcus thermophilus* (Sth1Cas9), *Neisseria meningitidis* (Nme1Cas9 and Nme2Cas9), *Lachnospiraceae bacterium (*LbaCas12a), and *Acidaminococcus sp*. (AspCas12a). We did a side-by-side comparison of editing efficiencies between the various systems using our simple assay. The gene encoding each CRISPR enzyme was cloned into a common vector for *in vitro* transcription of mRNA. For each system, we designed an assay for generating sgRNAs or crRNAs using PCR extension of annealed DNA oligos followed by *in vitro* transcription. Next, we performed the mutagenesis assay described previously. Following a microinjection of CRISPR-Cas messenger RNA (mRNA) and a pool of sgRNAs/crRNAs, fish were screened for pigmentation loss (**Figure 2A,B**). This screen demonstrated that SpyCas9 and SauCas9 were functioning relatively efficiently, as expected (**Figure S1**). However, injection of LbaCas12a, Sth1Cas9, or Nme1Cas9 as mRNAs, did not result in any pigmentation loss, suggesting no or inefficient editing of *tyr*.

We next performed a similar mutagenesis assay, utilizing commercially-available LbaCas12a enzyme instead of in vitro transcribed mRNA. We compared three forms of LbaCas12a crRNAs—chemically-synthetized processed crRNAs, in vitro transcribed pre-crRNAs, and in vitro transcribed processed crRNA with two guanines at the 5’ end for T7 transcription (Moreno-Mateos et al. 2017; P. Liu et al. 2019). When co-injected as LbaCas12a RNPs, all three forms of crRNA induced pigmentation loss (**Figure S2**). All forms of crRNAs exhibited increased activity when embryo growth temperature was increased to 34°C for four hours during early development, presumably when CRISPR activity is highest (Moreno-Mateos et al. 2017; P. Liu et al. 2019). We observed that in vitro synthesized pre-crRNAs and chemically-synthesized processed crRNAs were the most effective with ∼80% of embryos completely lacking pigmentation. Recently, an engineered version of AspCas12a, termed AsCas12a Ultra, was described with higher activity at a broad range of temperatures (Zhang et al. 2021). It recognizes the same PAM as LbaCas12a but with a distinct crRNA sequence. We tested the activity of AspCas12a Ultra (also known as AsCas12a Ultra) in targeting the *tyr* gene using in vitro synthesized pre-crRNAs and found that it was highly effective at both 28°C and 34°C (**Figure S2**). We concluded that LbaCas12a and AspCas12a are commercially-available CRISPR systems with high activity in zebrafish embryos.

By contrast, Sth1Cas9 mRNA and Nme1Cas9 mRNA were expressed but were not effective in inducing *tyr* mutant phenotypes (**Figure S1**). We were unable to obtain Sth1Cas9 or Nme1Cas9 in enzyme form. We attempted to troubleshoot their activity for mRNA injection by re-synthesizing sgRNAs, using alternative sgRNA scaffolds, and growing injected embryos at a higher incubation temperature; however, none of these attempts were able to induce pigmentation loss. More recently, an ortholog of Nme1Cas9, named Nme2Cas9, was described with increased activity in mammalian cells and animals, with a distinct NNNNCC PAM (Edraki et al. 2019). Injection of Nme2Cas9 RNPs into zebrafish embryos induced mosaic pigmentation loss in a sensitized (*tyr* +/-) genetic background, while Sth1Cas9 mRNA / sgRNA injection did not (**Figure S3A,B**). We confirmed Nme2Cas9 activity, and the lack of indels generated by Sth1Cas9, using T7 endonuclease I assay (**Figure S3C**). Since these CRISPR systems have non-overlapping PAMs (**Figure 1**), we suggest that they provide an improved set of tools for expanded targeting of double-strand DNA breaks to specific sequences in genomes.

For our remaining experiments, we proceeded with commercially-available and highly-active SpyCas9, SauCas9, and LbaCas12a enzymes and pools of guide RNAs for all microinjections. SpyCas9, SauCas9, and LbaCas12a all proved highly effective in disrupting the *tyr* gene (**Figure 2C,D**). Injections of SpyCas9 RNPs resulted in 100% pigmentation loss in all embryos (19/19). Injections of SauCas9 RNPs resulted in complete loss of pigmentation in 59% of embryos (35/59) and nearly complete loss in another 35% of embryos (21/59). Injections of LbaCas12a RNPs resulted in complete loss of pigmentation in 80% of embryos (63/78) and nearly complete loss in an additional 16% (13/78). Overall, we observed efficient rates of homozygous mutations at the *tyr* gene for all three systems with direct injection of commercially-available Cas protein and pools of guide RNAs into zebrafish embryos.

Next, we tested whether these systems were orthogonal. We defined the term orthogonal to mean the CRISPR-Cas systems can only function by utilizing their corresponding sgRNAs/crRNAs and not the sgRNAs/crRNAs from other systems. We expected that all three systems would be fully orthogonal, given their evolutionary distance and distinct PAM motifs (Esvelt et al. 2013). To confirm this, we tested each of the three CRISPR-Cas systems with each of the three pools of sgRNAs/crRNAs targeting the *tyr* gene and screened for pigmentation at 2 dpf. As expected, each of the CRISPR-Cas systems was only functional with its corresponding sgRNAs or crRNAs (**Figure 2B,D**). When injected with the sgRNA or crRNA pool from a different system, there was no evidence of gene editing. This confirms that SpyCas9, SauCas9, and LbaCas12a are fully orthogonal.

### Multiplexed F0 mutagenesis with orthogonal CRISPR-Cas systems

While SpyCas9 has been widely adopted in zebrafish, SauCas9 and LbaCas12a have only been tested at a small number of genes. We tested the efficacy of these two systems in phenocopying other classic mutants beyond *tyr*. We selected four genes with distinct mutant phenotypes: the transcription regulators *tbxta, tbx16*, and *noto* (all involved in mesoderm patterning), and *rx3* (essential for eye formation) (Halpern et al. 1993; Schulte-Merker et al. 1994; Kimmel et al. 1989; Griffin et al. 1998; Talbot et al. 1995; Halpern et al. 1995; Amacher and Kimmel 1998; Loosli et al. 2003). We designed pools of guide RNAs for each gene for both SauCas9 and LbaCas12a. We found that both systems were effective in generating robust phenocopy of the stable mutants, with >50% of embryos exhibiting complete mutant phenotype in all cases, with minimal non-specific effects (**Figure 3A,B**). We conclude that similar to SpyCas9, SauCas9 and LbaCas12a have high activity when co-injected with pools of guide RNAs and they may be broadly useful for CRISPR screening in animals.

**Figure 3.**
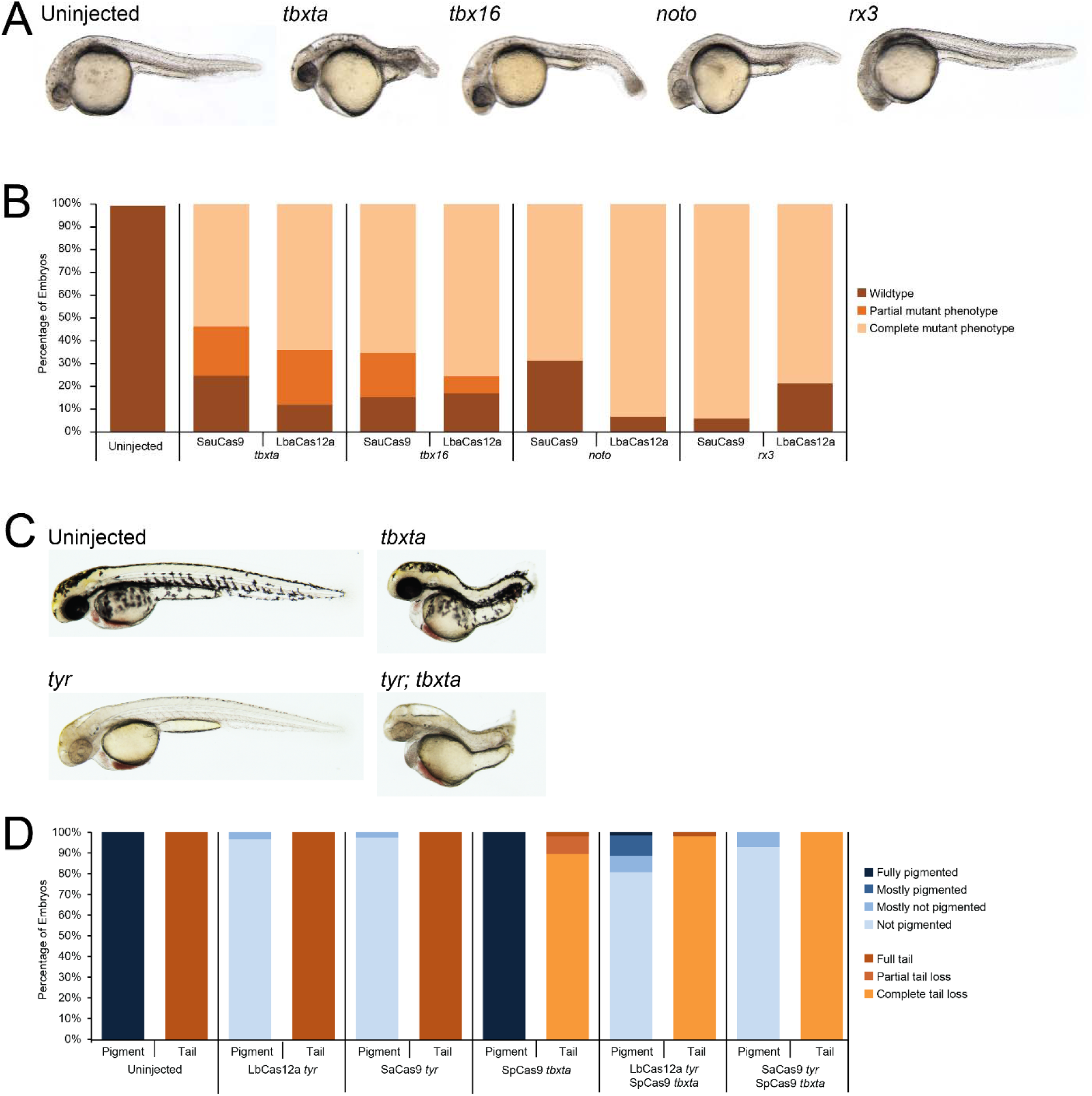
Multiplex mutagenesis with orthogonal CRISPR systems. **A**. Images of representative phenotypes for F0 mutagenesis of *tbxta, tbx16, noto*, and *rx3*. Embryos were injected with LbaCas12a and their corresponding guide RNAs at the 1-cell stage. Embryos were kept at 34C for the first four hours of development, then placed at 28C. Negligible toxicity was observed at 24 hpf. All embryos were imaged at 24 hpf. **B**. Quantification of embryonic phenotypes for mutagenesis of four genes, using both SauCas9 and LbaCas12a. The phenotypes were scored as wildtype, partial, or complete mutant phenotype for *tbxta* and *tbx16*, or just wildtype or complete mutant phenotype for *noto* and *rx3*, as these embryos did not exhibit obvious partial phenotypes. **C**. Images of representative embryos. Embryos representing uninjected and CRISPR-mutagenized embryos, as labeled, were imaged at 2 dpf, ideal for imaging both phenotypes. **D**. Quantification of embryonic pigmentation and tail phenotypes in categories described by the legend. Pigmentation phenotypes were categorized in the same way, as shown in Figure 2. Tail phenotypes were classified as full tail, partial tail loss, and complete tail loss. The latter is a complete phenocopy of the published *tbxta* mutant. Raw data in Table S3.

These results suggested that orthogonal CRISPR-Cas systems could be used simultaneously to disrupt multiple genes in the same individual. To test this, designed a set of SpyCas9 guide RNAs targeting *tbxta* and verified that injected embryos phenocopied the *tbxta* mutant with high penetrance (**Figure 3C,D**). Next, we generated double mutants using combinations of LbaCas12a, SauCas9 and SpyCas9 targeting *tyr* and *tbxta* simultaneously in the same embryos. We observed high rates of mutagenesis at both genes, with >90% of embryos exhibiting complete phenocopy of both mutants (**Figure 3C,D**). This experiment demonstrates that efficient multiplexed mutagenesis in zebrafish embryos is possible with orthogonal CRISPR-Cas systems.

### CRISPR recording using orthogonal CRISPR systems

Current CRISPR recording methods rely on SpyCas9. We hypothesized that orthogonal CRISPR systems could be used to improve CRISPR recording. We generated a stable transgenic zebrafish containing a single-copy array, termed a “barcode,” composed of 5 sites for each of three CRISPR systems. To deliver CRISPR edits to this barcode, we used 1-cell injection of SauCas9 and LbaCas9 RNPs and heat-shock induction of SpyCas9 and corresponding guide RNAs 24 hours after fertilization (**Figure 4A**). The next day, we isolated genomic DNA from individual embryos and used Illumina amplicon sequencing to measure barcodes. Uninjected embryos had no edits at barcodes, while delivery of single CRISPR systems induced edits only at the corresponding target sites at the barcode, as expected (**Figures 4B,S4**). Delivery of all three CRISPR systems induced robust editing across the barcode (**Figure 4B**). Individual animals edited with all three CRISPR systems contained thousands of unique barcodes, with ∼5 edits per barcode on average (**Figure 4C,D**). These barcodes contain more edits than previous GESTALT and scGESTALT barcodes (**Figure 4C**) (McKenna et al. 2016; Raj et al. 2018), which translates to more information density for lineage or molecular recording. Barcodes edited with all three CRISPR systems were highly diverse in comparison with barcodes edited with single systems (**Figures 4D,S5**). As these systems are truly orthogonal, they avoid issues with crosstalk between editing time points, which occurred at low incidence in scGESTALT multi-timepoint recording.

**Figure 4.**
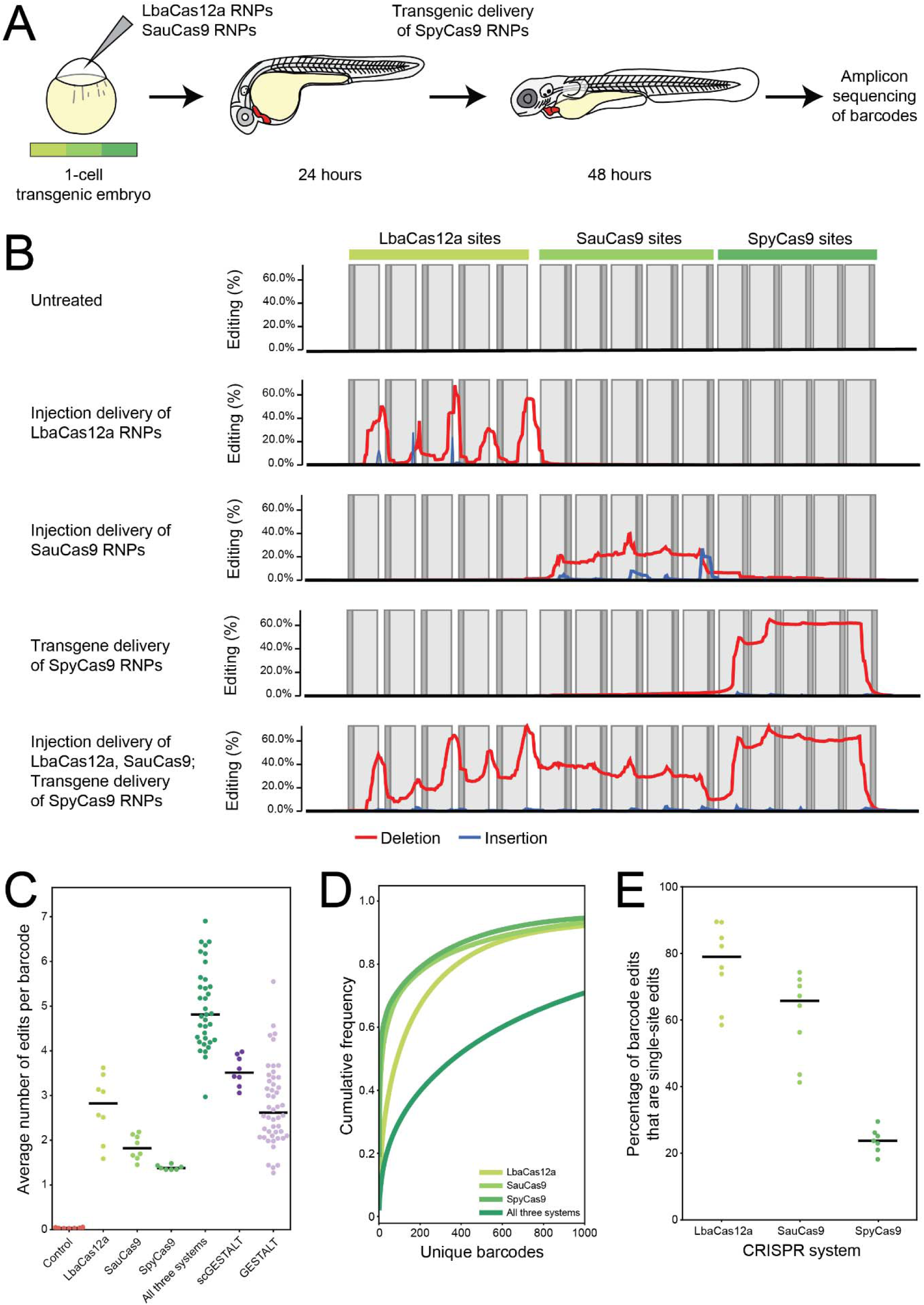
CRISPR recording with orthogonal CRISPR systems. **A**. A diagram of the experimental design. **B**. Barcode edit plots display the mutations within the 15 CRISPR target sites (5 for each CRISPR system) of the barcode for five editing conditions. Percentage of barcodes containing an edit at each nucleotide position is shown. One representative animal is shown per condition; additional plots can be found in **Figure S4**. The PAM is shown in dark grey, target site in light grey. Red lines represent deletions; blue lines represent insertions. **C**. Average number of edits per barcode, weighted by the barcode abundance in a given sample, is shown for each condition. scGESTALT and GESTALT data were from previous publications (McKenna et al. 2016; Raj et al. 2018). Each dot represents an individual animal. Lines indicate the median within each group. LbaCas12a generates more edits, on average, than SauCas9 (*p\** = 0.005415) and SpyCas9 (*p*\* = 0.0006348). Our system combining LbaCas12a, SauCas9, and SpyCas9 produces more edits in a barcode than scGESTALT (*p*\* = 1.492 × 10^−8^) and GESTALT systems (*p*\* < 2.2 × 10^−16^). **D**. Cumulative frequency of barcodes from within a single representative animal for each condition. The legend identifies each sample. **E**. The percentage of total edits that were single-site edits (local edits/total edits) is shown for each CRISPR system. Each dot represents the weighted average of all barcodes sequenced from a single animal; lines indicate the median within each group. LbaCas12a generates more single-site edits than SauCas9 (*p*\* = 0.01227) and SpyCas9 (*p*\* = 7.319 × 10^−7^).

Finally, we observed that LbaCas12a CRISPR editing of barcodes resulted in a bias towards single-site edits, rather than the multi-site edits observed with SauCas9 and SpyCas9 (**Figure 4B,E**). ∼80% of the edits generated by LbaCas12a were single-site edits. Multi-site edits have two distinct negative consequences—first, they occupy multiple unedited sites, reducing information density, and second, they may erase previous edits, leading to errors in lineage reconstruction or molecular recording (Salvador-Martínez et al. 2019). We hypothesize that the staggered breaks induced by Cas12a systems have different repair kinetics, biasing outcomes to single-site edits that are more favorable for confident CRISPR recording (Yeh, Richardson, and Corn 2019). In any case, we suggest that orthogonal CRISPR systems offer improved tools for high-density, high-diversity multichannel CRISPR lineage tracing and molecular recording.

### Anti-CRISPR proteins can efficiently block CRISPR-Cas genome editing in zebrafish embryos

We hypothesized that anti-CRISPR proteins - small peptides that block the activity of Cas enzymes - would be effective inhibitors of CRISPR-Cas activity in zebrafish. We selected four type II Acr proteins that had previously been proven functional in bacterial and human cells: AcrIIA2, AcrIIA4, AcrIIC1, and AcrIIC3 (Pawluk et al. 2016; Rauch et al. 2017). We cloned each into a common vector for mRNA transcription. To test each Acr protein, they were co-injected as mRNAs alongside each CRISPR-Cas protein and its corresponding sgRNAs/crRNAs targeting *tyr*. Pigmentation was screened at 2 dpf (**Figure 5A**). We hypothesized that if the Acr was functional, it would block CRISPR-Cas editing of the *tyr* gene, resulting in the rescue of pigmentation. When injected with SpyCas9, AcrIIA2 and AcrIIA4 blocked editing with a high efficiency (**Figure 5B,C**). As expected, AcrIIC1 and AcrIIC3, previously found to inhibit type II-C CRISPR-Cas systems, had little inhibitory effect on editing. When injected with SauCas9, only AcrIIA4 induced a moderate level of inhibition on editing, with 20% of the embryos fully pigmented. As expected, none of the Acr proteins were effective in blocking editing when co-injected with LbaCas12a, as it is a type V CRISPR-Cas system. We did not test the ability of Acrs to inhibit Nme2Cas9, given the low activity of this CRISPR system in zebrafish embryos.

**Figure 5:**
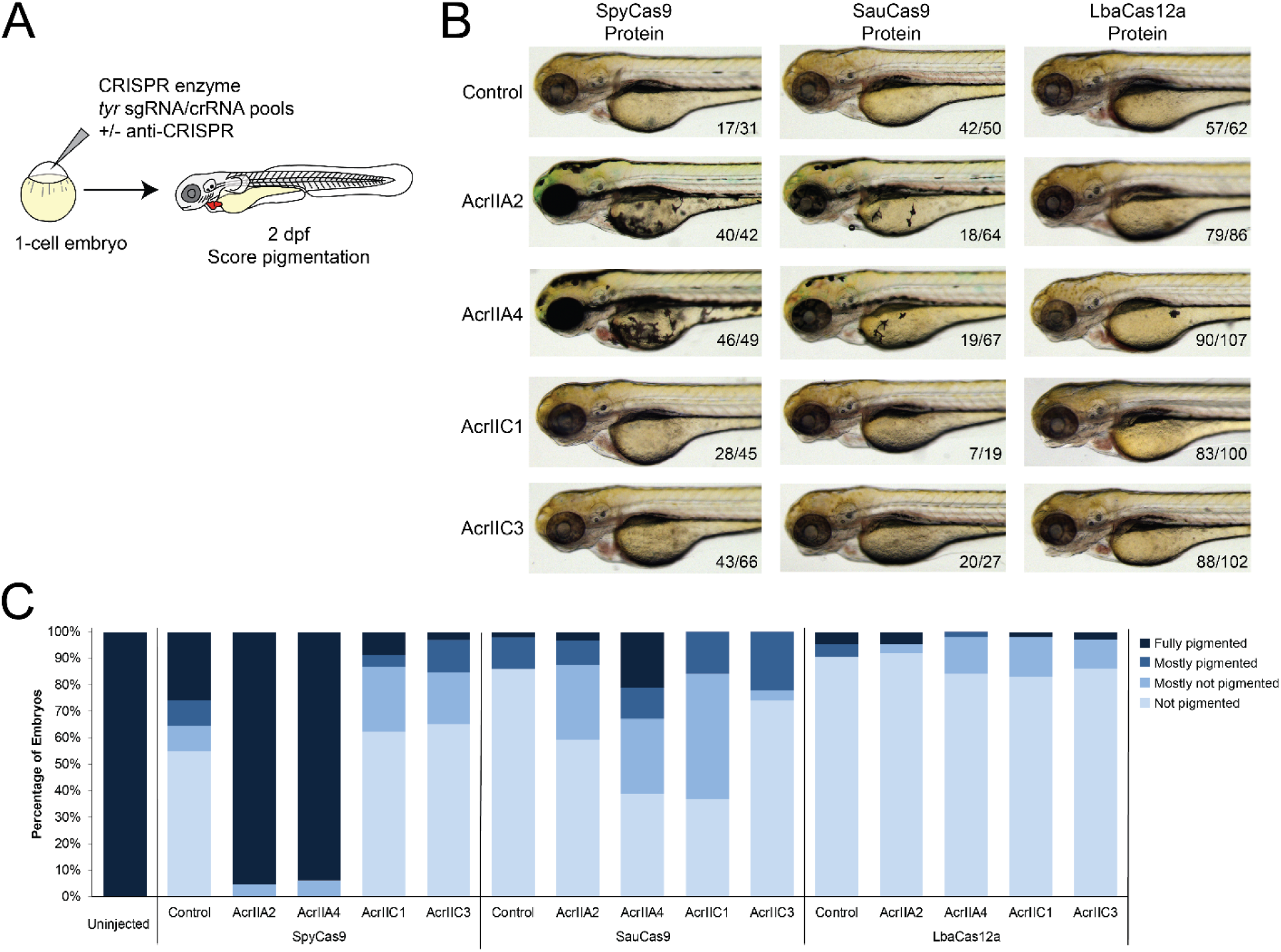
The activity of various anti-CRISPR proteins to inhibit SpyCas9, SauCas9, and LbaCas12a in zebrafish embryos. **A**. Experimental design of the assay for anti-CRISPR activity. A mix of CRISPR-Cas enzyme, *tyr* sgRNAs/crRNAs, and anti-CRISPR mRNA or water was microinjected into the single-cell zebrafish embryo. At 2 dpf the fish are screened for their level of pigmentation. **B**. Example images of embryos injected as described in panel A, for three CRISPR systems and four anti-CRISPR proteins. **C**. Quantification of pigmentation categories after microinjection as described. Raw data in Table S3.

We were initially surprised that anti-CRISPRs were so effective, since they were injected as mRNAs and needed to be translated in the embryo before they would be able to interfere with injected Cas protein activity. We hypothesized that Acr proteins would be more effective at inhibiting gene editing when co-injected with Cas9 mRNA. However, we found no difference in the level of CRISPR-Cas inhibition between injected SauCas9 mRNA or protein **(Figure S6**). We conclude that AcrIIA2 and AcrIIA4 are highly effective inhibitors of SpyCas9, with moderate activity against SauCas9. More CRISPR inhibitor proteins have recently been described, and our simple and rapid assay for gene editing may serve as a platform for further screening for effective inhibitors of other CRISPR systems in zebrafish.

## Discussion

In this study, we described a simple CRISPR-Cas mutagenesis screen and demonstrated highly-efficient activity of three orthogonal CRISPR-systems in zebrafish: SpyCas9, SauCas9, and LbaCas12a. We also demonstrated high activity of AspCas12a, which has an overlapping PAM with LbaCas12a, and moderate genome editing in zebrafish using RNP delivery of Nme2Cas9. We conclude that the lesser-used SauCas9 and LbaCas12a systems are similarly potent to SpyCas9 when delivered as RNPs and can be used as effective tools for F0 mutagenesis. In our hands, LbaCas12a was only functional when delivered as an RNP, with either synthetic or in vitro transcribed crRNAs. It is possible that direct RNP injection of Nme1Cas9 or Sth1Cas9 could also rescue activity. We also showed that Acr proteins can be used for CRISPR-Cas inhibition in zebrafish. AcrIIA2 and AcrIIA4 were both effective inhibitors of SpyCas9, and to a lesser extent, AcrIIA4 inhibited SauCas9 activity. AcrIIC1 and AcrIIC3 have previously been shown to inhibit CRISPR-Cas activity in type II-C systems, so it is not surprising they were not effective against SpyCas9, SauCas9, and LbaCas12a. As exploration of different types of CRISPR-Cas systems and Acr proteins continue, their use for CRISPR regulation will grow.

Our findings expand the toolkit of CRISPR-Cas systems in zebrafish and demonstrate some initial intersectional applications. We anticipate several areas in which simultaneous or intersectional applications of these systems could be beneficial. First, since each CRISPR-Cas system utilizes a unique PAM, more regions of the genome are now available for mutagenesis. Second, as LbaCas12a and AspCas12a generate staggered DSBs, this system may generate distinct repair outcomes, or be more likely to drive homology-directed insertion events (Moreno-Mateos et al. 2017; P. Liu et al. 2019), with potential advantages over SpyCas9 or SauCas9 for certain applications. Third, the use of orthogonal CRISPR-Cas systems in the same individual opens the door for combining CRISPR modalities for more sophisticated screens, for example, combining F0 mutagenesis with simultaneous CRISPR activation, repression, base editing, or epigenetic modification (K. Liu et al. 2019; Thakore et al. 2016). Fourth, anti-CRISPR proteins offer a new strategy for regulated CRISPR activity in specific tissues or at particular developmental times, through the use of tissue-specific or inducible promoters. This spatiotemporal control will also enable more sophisticated genetic approaches.

We found that orthogonal CRISPR systems can enable more sophisticated CRISPR recording tools. We show that barcodes edited with three CRISPR systems contain more edits than previous methods. We also show that LbaCas12a, which has not been previously used for CRISPR recording, biases DNA repair outcomes in favor of single-site edits instead of the multi-site edits common with SpyCas9 and SauCas9 barcode editing. This will increase information density and reduce issues with erased recordings. We anticipate that orthogonal systems will also enable recording at more timepoints in the same individual, leading to higher confidence in the resulting lineage trees. There is similar potential for multi-channel recording with distinct CRISPR systems, for example, by using one system to record lineage and another to record cell signaling. While less effective for F0 mutagenesis, Nme2Cas9 could be useful for longitudinal CRISPR recording where the slower accumulation of edits would be a valuable feature, especially considering the lower propensity of Nme2Cas9 for off-target editing (Edraki et al. 2019). We imagine that anti-CRISPR proteins may serve a similar role in dampening recording intensity in favor of recording over longer durations of time.

Together, our study supports the continual exploration and application of emerging genome editing tools in zebrafish. The zebrafish offers immense practical advantages over nearly all other vertebrate model organisms as a testbed for genome engineering. Here, we have shown the functionality of several CRISPR-Cas systems and Acr proteins in zebrafish and repurposed them for CRISPR recording. It is likely that many more CRISPR-Cas and anti-CRISPR systems can be ported to zebrafish, offering further opportunities for targeting and multiplexing. We anticipate that sophisticated genetic strategies enabled by multiplexed CRISPR systems will have a large impact on how we study vertebrate development and model human diseases in other animals.

## Materials and Methods

### Zebrafish husbandry

All vertebrate animal work was performed at the facilities of the University of Utah, CBRZ. This study was approved by the Office of Institutional Animal Care & Use Committee (IACUC # 18-2008) of the University of Utah’s animal care and use program.

### Cloning and transcription of CRISPR-Cas systems and anti-CRISPR

CRISPR-Cas and anti-CRISPR plasmids were ordered from Addgene, thanks to gifts from many investigators (Hou et al. 2013; Gagnon et al. 2014; Ran et al. 2015; Kleinstiver et al. 2015; Pawluk et al. 2016; Moreno-Mateos et al. 2017; Rauch et al. 2017). Open reading frames were amplified using PCR with primers that add overlapping ends corresponding to the pCS2 vector, and subcloned into the pCS2 vector using NEBuilder HiFi DNA Assembly (NEB). All oligo sequences are available in **Table S2**. Plasmids were miniprepped (Zymo) and confirmed via Sanger sequencing. Each vector was linearized using NotI restriction digest (NEB). Capped mRNA was synthesized using the HiScribe SP6 kit (NEB) and purified using the RNA Clean & Concentrator kit (Zymo). All constructs generated in this study are available at https://www.addgene.org/James_Gagnon/.

### Generation of sgRNAs and crRNAs

SpyCas9 *tyr* and *tbxta* sgRNAs were synthesized using EnGen sgRNA Synthesis Kit (NEB). SauCas9, Sth1Cas9, and Nme1Cas9 sgRNAs were synthesized as previously described, with modifications (Gagnon et al. 2014). Briefly, gene-specific and constant oligos were designed for overlap extension for template synthesis. A reaction containing 2 ul constant oligo (5 uM), 2 ul gene-specific oligo (5 uM), 12.5 ul 2X Hotstart Taq mix, and 8.5 ul water was cycled on a thermocycler using this protocol - 95°C for 3 mins, then 30 cycles of (95°C 30 seconds, 45°C 30 seconds, 68°C 20 seconds), followed by 68°C for 5 minutes. Templates were run on a 1% TAE agarose gel to confirm correct band size, and purified using the DNA Clean and Concentrator kit (Zymo). sgRNAs or crRNAs were transcribed using the HiScribe T7 High Yield RNA Synthesis kit (NEB) or the MEGAscript T7 Transcription kit (Thermo Fisher), and purified using the RNA Clean & Concentrator kit (Zymo), or by using phenol chloroform RNA extraction. AspCas12a and LbaCas12a *tyr* crRNAs were transcribed following a previously-described protocol (P. Liu et al. 2019), or chemically synthesized (Synthego). Nme2Cas9 sgRNAs were generated using a previously published protocol (Amrani et al. 2018; Edraki et al. 2019). The sgRNAs or crRNAs were then pooled into a single mix at equal molarities (∼600 ng/ul for sgRNAs, ∼1800 ng/ul for crRNAs). For SpyCas9 we pooled 4 or 5 sgRNAs, SauCas9 we pooled 5 sgRNAs, Nme1Cas9 we pooled 10 sgRNAs, Nme2Cas9 we pooled 10 sgRNAs, Sth1Cas9 we pooled 3 sgRNAs, and LbaCas12a we pooled 3 crRNAs into the final pools we used to make injection mixes. All oligo sequences are in **Table S2**.

### Cas9 CRISPR-Cas injection mixes

We assembled microinjection mixes in the following order in 1.5 ml tubes: 1 ul of 1 M KCl (Sigma-Aldrich), 0.5 ul phenol red (Sigma-Aldrich), 1 ul of a mix of sgRNAs, generated as described above. This pre-mix was briefly vortexed and centrifuged to bring the solution to the bottom of the tube. Then 1 ul Cas mRNA (∼300 ng/ul), 1 ul of 20 μM Cas protein (SpyCas9 and SauCas9 from NEB; Nme2Cas9 was purified as previously described (Edraki et al. 2019)), and/or 1 ul of anti-CRISPR mRNA (∼500 ng/ul) was added to the tube, and the mix vortexed and centrifuged again. Nme2Cas9 injection mixes were incubated at 37C for 5 minutes, then kept on ice until ready. 1-2 nl was injected into the cell of a zebrafish zygote. Nme2Cas9 and Sth1Cas9 were injected into tyr +/- embryos for more sensitive mutation detection, and wildtype embryos for T7E1 assay, as shown in **Figure S3**.

### Cas12a CRISPR-Cas injection mixes

We assembled microinjection mixes in the following order in 1.5 ml tubes: 1 ul of 1 M KCl (Sigma-Aldrich), 0.5 ul phenol red (Sigma-Aldrich), 1.5 ul (LbaCas12a) or 2.5 ul (AspCas12a) of a mix of crRNAs, generated as described above. This pre-mix was briefly vortexed and centrifuged to bring the solution to the bottom of the tube. Then 1.5 ul of 50 μM LbaCas12a protein (NEB) or 1 ul of 63 μM AsCas12a Ultra protein (IDT) was added to the tube, and the mix vortexed and centrifuged again. Injection mixes were incubated at 37C for 5 minutes, then kept on ice until ready. 1-2 nl was injected into the cell of a zebrafish zygote. Injection mixes for multiplexed mutagenesis were generated by doubling the pre-mix and Cas protein volumes.

### CRISPR mutagenesis assays

At 1 dpf, we screened to remove unfertilized or dead embryos. In most conditions, <10% of embryos exhibited toxicity, which we attribute mostly to injection artifact. At 2-3 dpf, embryos were scored for pigmentation loss into one of four categories: fully pigmented (100% pigmentation), mostly pigmented (51-99% pigmentation), mostly not pigmented (6-50% pigmentation), and not pigmented (0-5% pigmentation) (**Figure 2B**). T7 endonuclease 1 assay was performed following manufacturer’s protocol (NEB).

### Imaging

Larvae were imaged at 3 dpf, with the exception of the embryos in **Figure 3**, which were imaged at 1 or 2 dpf. In cases where the larvae had not hatched from the chorion, they were manually dechorionated using tweezers. From a stock solution of 4 mg/ml of Tricaine (Sigma-Aldrich), we create a diluted solution of 0.0064 mg/ml in E3 buffer. The larvae were anesthetized in diluted Tricaine solution for 2 minutes. Once the larvae were immobile, they were moved onto a thin layer of 3% methylcellulose (Sigma-Aldrich) and oriented for a lateral or top view. Images of larvae were taken using a Leica M205FCA microscope with a Leica DFC7000T digital camera.

### Barcode design

Well-edited CRISPR targets that were not present in the zebrafish genome were identified for SauCas9 and LbaCas12a using a combination of computational design, literature review, and experimental validation. SpyCas9 targets were used from previous barcode designs (McKenna et al. 2016). Five sites for each of the three systems were concatenated in an array, with a three nucleotide spacer between each target site, and cloned into a vector containing a *myl7*:GFP marker and Tol2 recognition sites for transgenesis. This transgenesis vector was named pTol2-HybridBarcode, and is available at https://www.addgene.org/James_Gagnon/.

### Generation of transgenic zebrafish with a single-copy CRISPR barcode

To generate founder fish, 1-cell embryos were injected with Tol2 mRNA and pTol2-HybridBarcode vector DNA. Potential founder fish were screened for GFP expression in the heart at 30 hpf and grown to adulthood. Founder transgenic fish were identified by outcrossing to wild type and screening clutches of embryos for heart GFP expression at 30 hpf. A single-copy Tol2 transgenic line was identified from a single founder using copy number qPCR as previously described (McKenna et al. 2016). This barcode line was given the ZFIN line designation *zj1Tg*, expanded by out-crossing, and used for all experiments in this manuscript.

### CRISPR barcoding

The barcode line^*zj1Tg*^ was crossed to wild type or Tg(hsp70l:zCas9-2A-EGFP,5x(U6:sgRNA))^a168Tg^ males (Raj et al. 2018) to generate double transgenic embryos. SauCas9 and/or LbaCas12a RNPs containing the appropriate guide RNAs were injected at the 1-cell stage with 2 nl of the following injection mix: 0.75 ul 1M KCl, 1.75 ul sgRNA/crRNA mix at molar ratios, 1.25 ul SauCas9 protein, 0.75 ul LbaCas12a protein, 0.5 ul phenol red. These embryos were screened for green hearts to identify the barcode transgene, and then heat shocked to induce SpyCas9 expression, as previously described (Raj et al. 2018), and grown to 2 days of age. Genomic DNA was extracted from individual embryos following the HotSHOT method.

### Amplicon sequencing of barcodes

Amplicon sequencing libraries were prepped using two rounds of PCR, which completed the Illumina adapters and added dual 8bp indices that were unique to each sample, following previously published protocols (Gagnon et al. 2014; Komor et al. 2016). Libraries were pooled at roughly equimolar ratios, and sequenced on an Illumina MiSeq using 600-cycle v3 kits.

### Computational analysis

Sequencing data was processed using the previously published GESTALT pipeline, with modifications to permit the use of Singularity as a container environment on our computing cluster. All code for post-processing and analyzing barcodes will be available at https://github.com/Gagnon-lab/.

### Statistical analysis

All statistical tests performed were unpaired, two-sample, one-tailed Student t-tests. The Welch t-test provided by the “t.test” function in R was used to calculate p-values in Figure 4C,E.

## Acknowledgements

We thank CZAR and CBRZ staff, especially Nathan Baker, for excellent zebrafish care. We thank all members of the Gagnon lab, Yannick Doyon, James Keener, Alexis Komor, Nathan Lawson, Aaron McKenna, and Michael Werner for their support, feedback, and suggestions on this project. We thank Addgene and the labs who contributed vectors for their commitment to open science. Support for this project came from NIH grant R35GM142950 (JAG), UROP grants from the University of Utah Office for Undergraduate Research (PRT, SNK), from the NSF Graduate Research Fellowships Program (KTT), from the ACCESS Program (SNK), from the Beckman Scholars Program (SNK), from the Biology Research Scholars Program (SNK), from the Ryan Watts Research Fellowship program (MAH), and from the Office of the Vice President for Research, the Henry Eyring Center for Cell & Genome Science, the 1U4U program, and startup funds from the University of Utah (JAG). We thank the Taft-Nicholson Center for space and support to write this manuscript. This research was conducted on the traditional and ancestral homeland of the Shoshone, Paiute, Goshute, and Ute Tribes. We affirm and support the University of Utah’s partnership with Native Nations and Urban Indian communities.

## Author contributions

PRT led the CRISPR and Acr screens, generated and validated the CRISPR barcoding animals, and led the writing of the manuscript. SW contributed to the LbaCas12a and AspCas12a studies. SW and KTT conducted and analyzed the CRISPR barcoding experiments. EPD, SNH, and MAH assisted with cloning, multiplex mutagenesis, and various experiments. NB, OO, ES expressed and purified Nme2Cas9, and contributed expertise with guide RNA design. JAG conceived of and supervised the project, assisted with experiments, and assisted with writing the manuscript.

## Data availability statement

The embryo phenotype scoring data is available in Table S3. Sequencing data is pending on GEO accession number. Vectors are available at https://www.addgene.org/James_Gagnon/. All code will be available at https://github.com/Gagnon-lab/.

## Supplementary information

**Figure S1.**
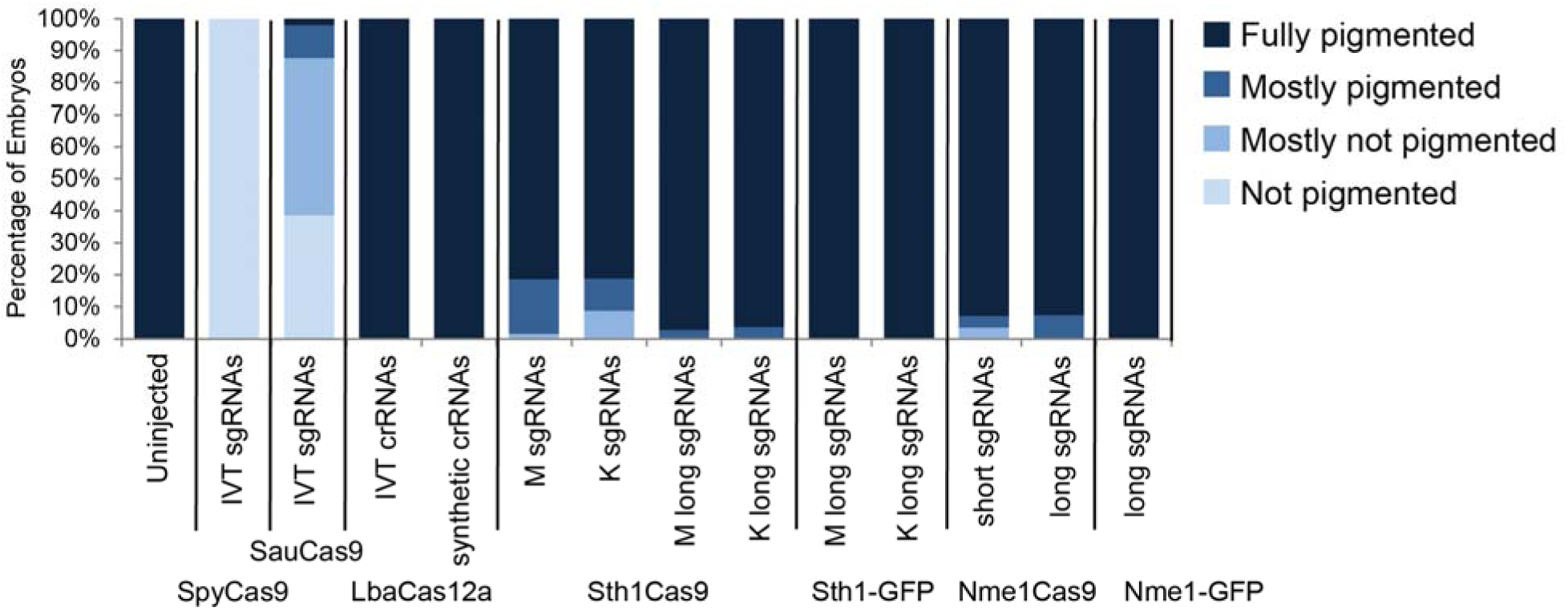
A screen of CRISPR systems in zebrafish embryos using mRNA injection and a variety of sgRNA and crRNA designs. Percentage of embryos exhibiting pigmentation loss at 2 dpf after targeting the *tyr* gene with pools of sgRNAs or crRNAs and mRNA encoding each Cas enzyme. LbaCas12a was tested with both in vitro transcribed crRNAs and synthetic crRNAs. Sth1Cas9 was tested with the sgRNA scaffolding from both Muller (M, Müller et al. 2016) or Kleinstiver (K, Kleinstiver et al. 2015), a lengthened sgRNA scaffold (long, Steinert et al. 2015) described in Steinert and Sth1Cas9-t2a-GFP mRNA construct. Similarly, Nme1Cas9 was tested with a lengthened (long) sgRNA scaffold and the Nme1Cas9-t2a-GFP mRNA construct.

**Figure S2.**
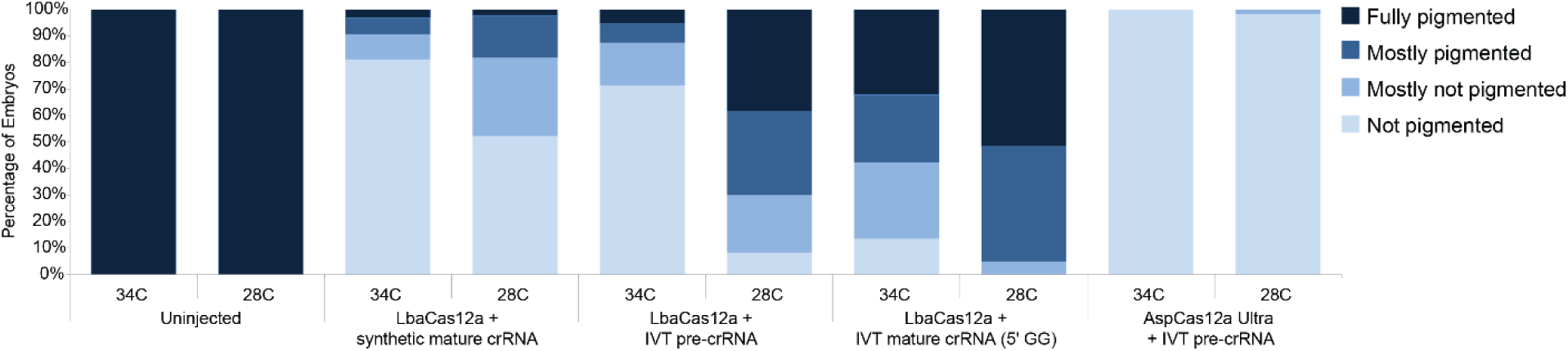
CRISPR mutagenesis with LbaCas12a and AspCas12a Ultra. Percentage of zebrafish embryos exhibiting pigmentation loss at 2 dpf after targeting the *tyr* gene with LbaCas12a or AspCas12a Ultra RNPs, RNPs with either synthetic mature (processed) crRNAs, in vitro transcribed pre-crRNAs, or in vitro transcribed mature crRNAs with a 5’GG appended to permit T7 transcription. Each injection mix was assembled to contain the same crRNA molarity. A pool of embryos was injected, and then split and incubated at either 28C or 34C for 4 hours post injection, then placed at 28C.

**Figure S3.**
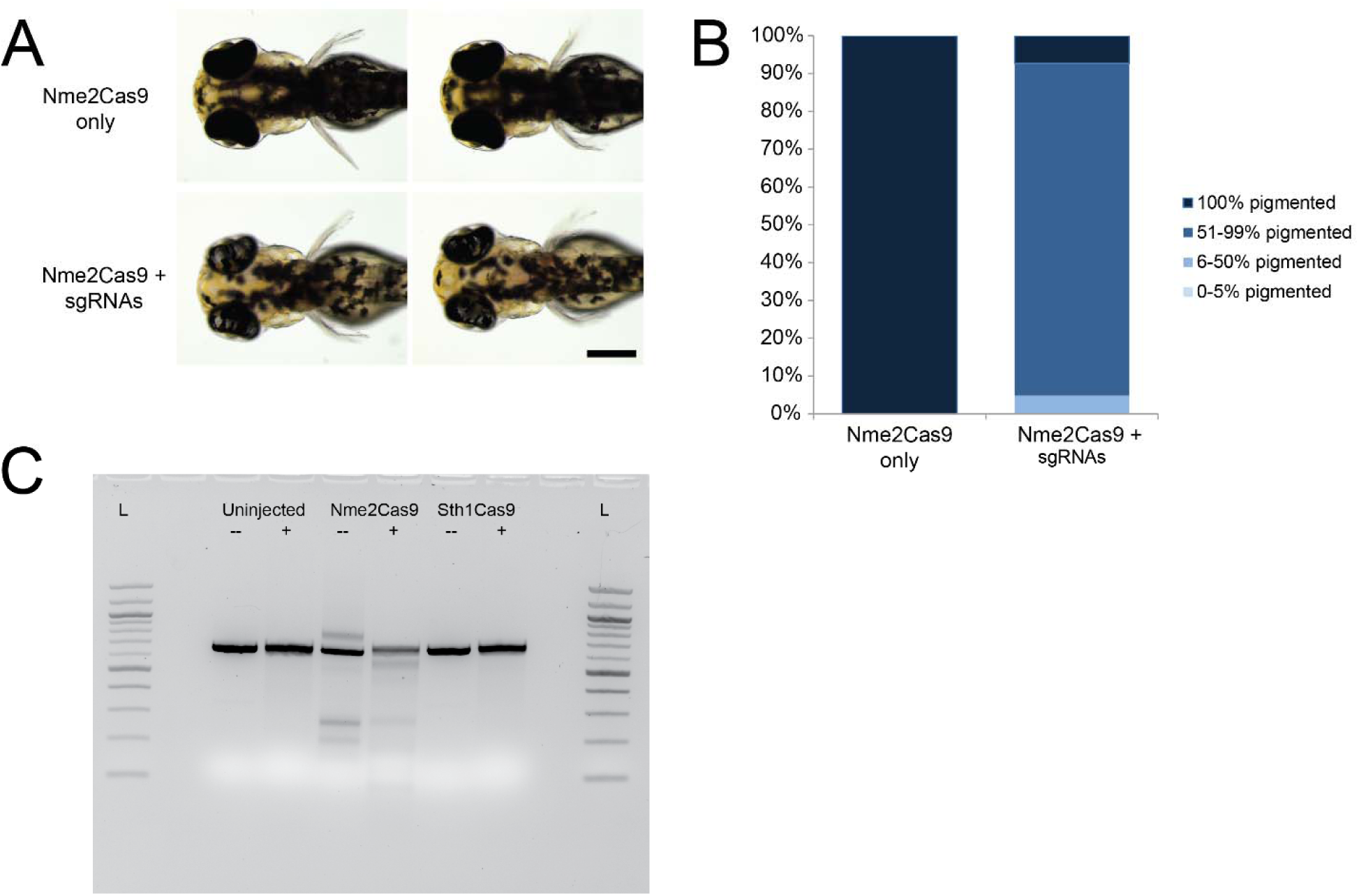
CRISPR editing of zebrafish embryos with Nme2Cas9. **A**. *tyr* +/- embryos injected with Nme2Cas9 RNPs targeting the *tyr* gene, or Nme2Cas9 alone, are shown at 3 dpf. Two representative embryos are shown for each condition. **B**. Percentage of injected zebrafish embryos exhibiting pigmentation loss at 3 dpf. 25-40% of embryos from both conditions exhibited deformities at 24 hpf and were removed before scoring. **C**. T7 endonuclease I assay on PCR product flanking the *tyr* target sites from uninjected wild type embryos, embryos injected with Nme2Cas9 RNPs, and embryos injected with Sth1Cas9 mRNA + sgRNAs. (-) indicate PCR product before T7 endonuclease I assay (expected size of 754bp), (+) indicates after T7 endonuclease I assay. Note that large deletions can be observed in the Nme2Cas9 RNP injected condition, even before T7 endonuclease I assay. These can be attributed to simultaneous activity of two or more sgRNAs.

**Figure S4.**
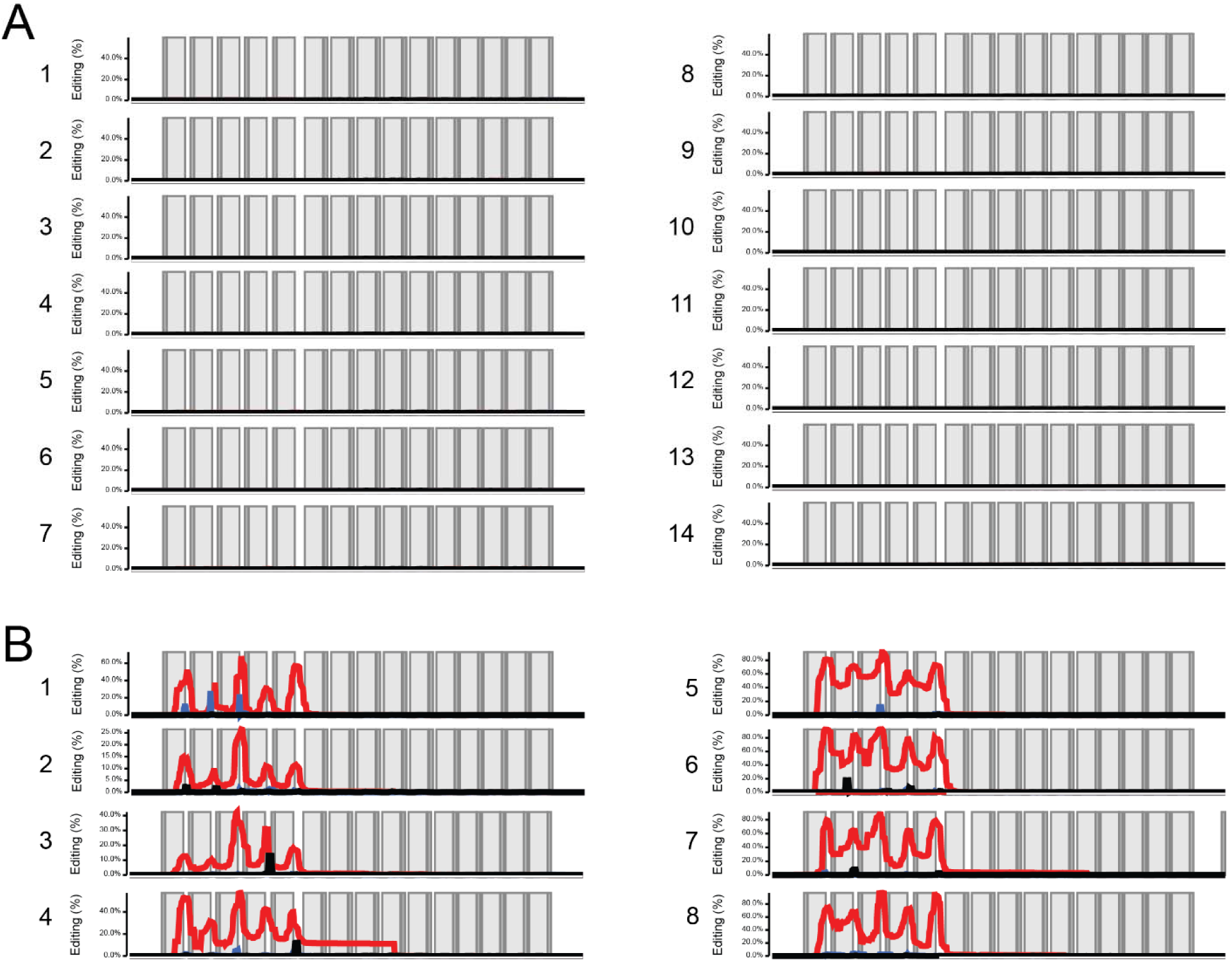
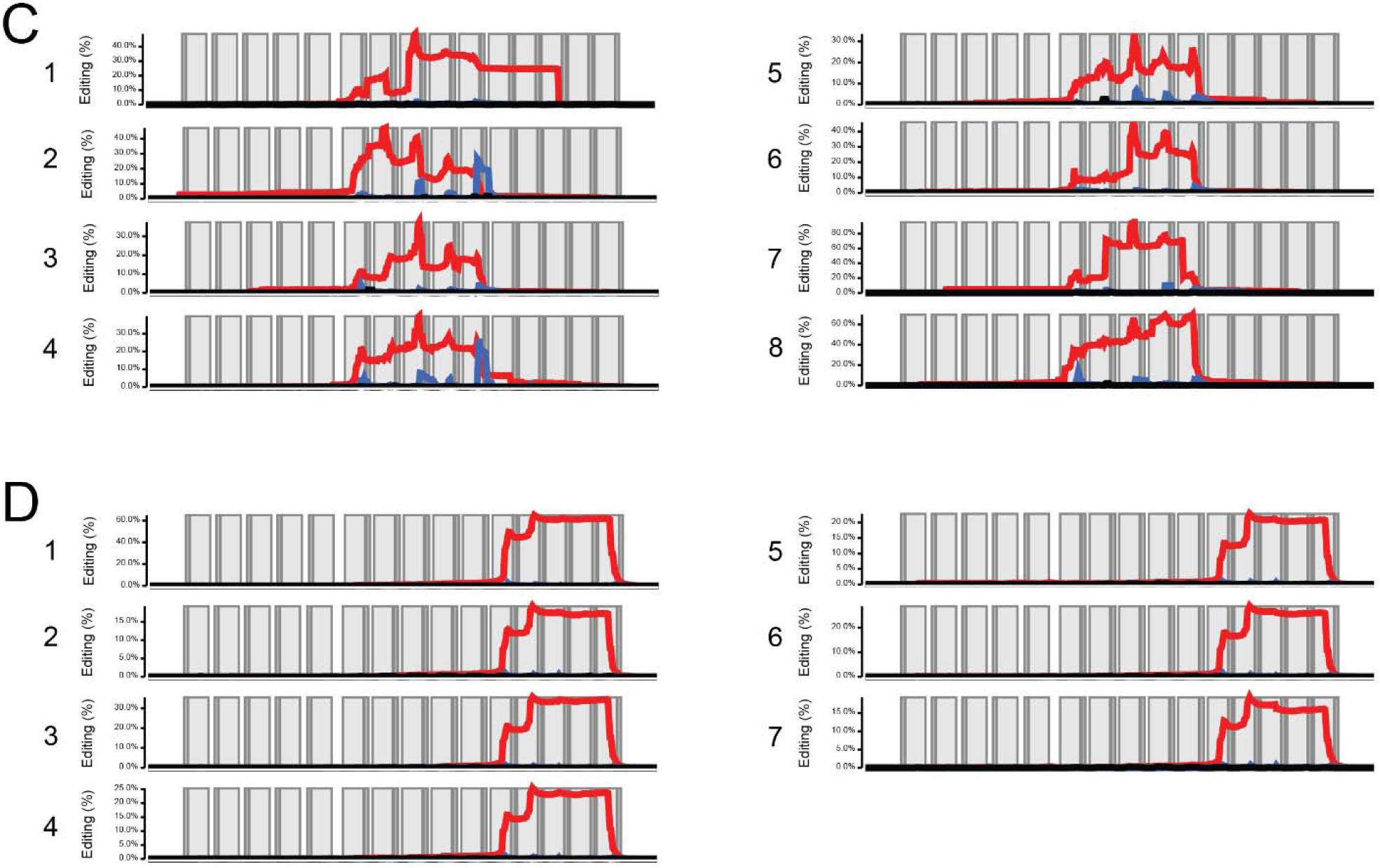
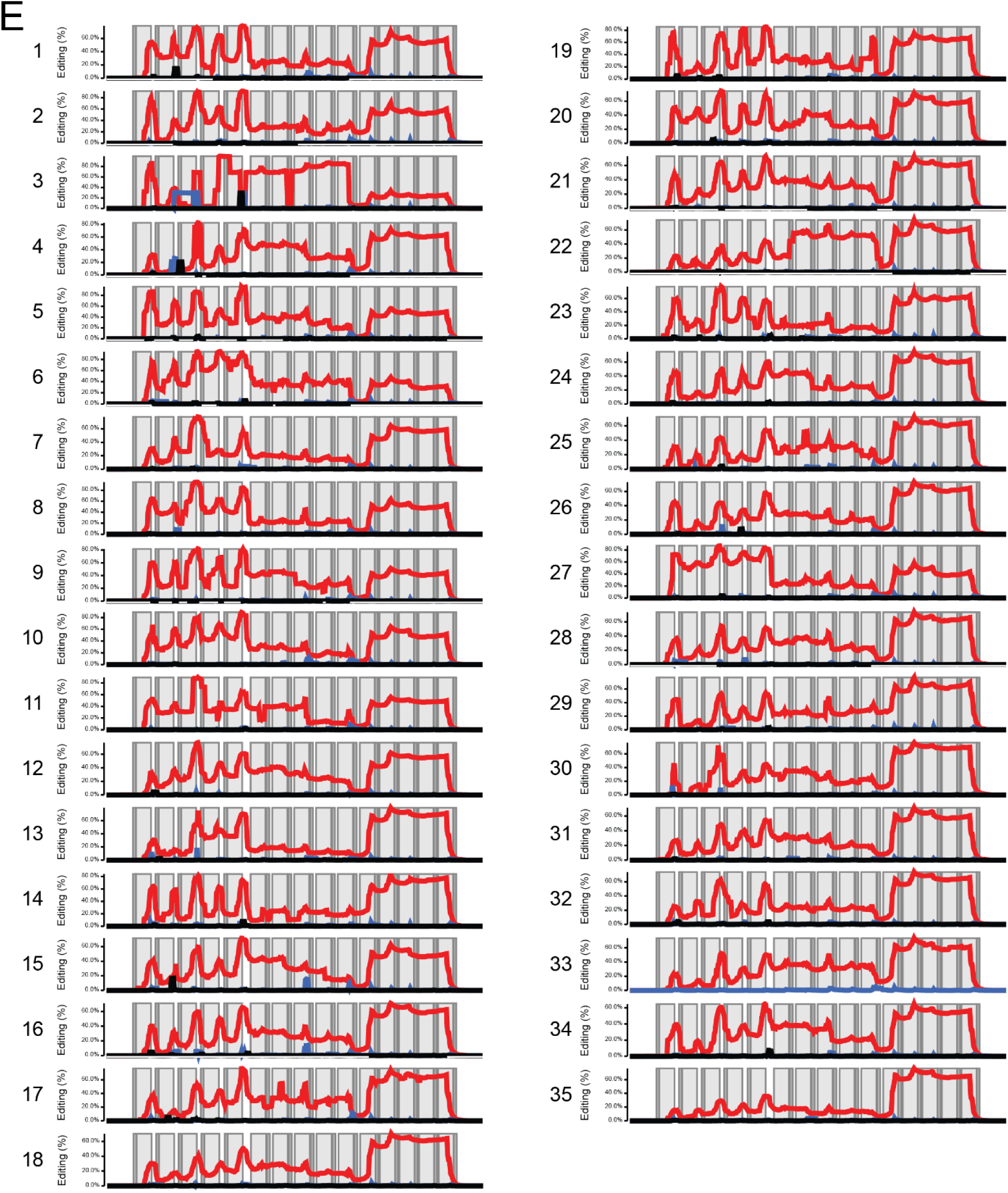
Barcode edit plots from individual embryos edited with orthogonal CRISPR systems. Each plot displays the mutations within the 15 CRISPR target sites (5 for each CRISPR system) of the GESTALT barcode from an individual embryo. The PAM for each CRISPR systems is shown in dark grey, target site in light grey. Red lines represent deletions, blue lines represent insertions, Black lines represent substitutions. **A**. Uninjected control embryos (n=14). **B**. Embryos edited by LbaCas12a RNP injection (n=8). **C**. Embryos edited by SauCas9 RNP injection (n=8). **D**. Embryos edited by induced transgenic delivery of SpyCas9 RNPs (n=7). **E**. Embryos edited with all three CRISPR systems (n=35).

**Figure S5.**
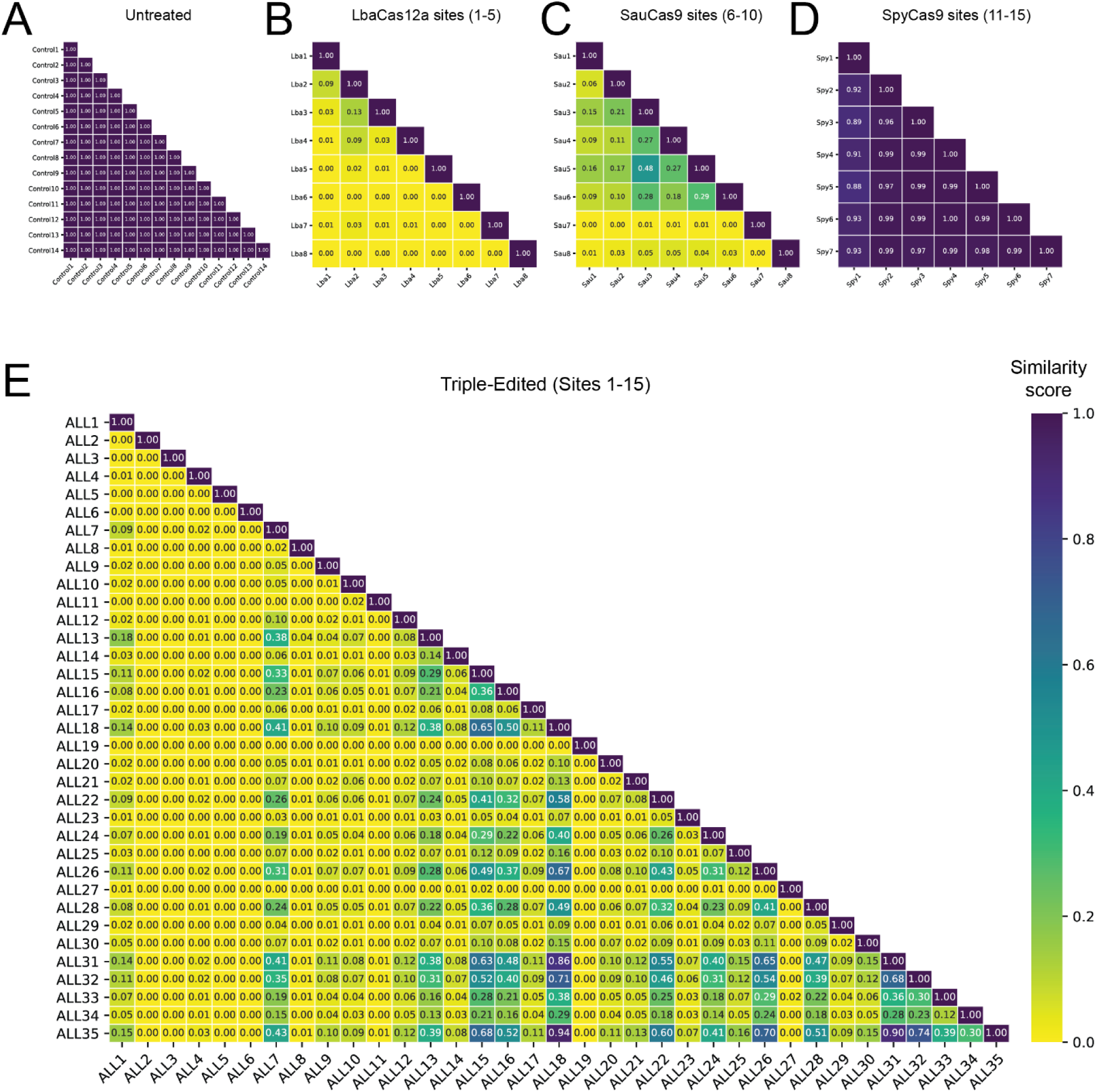
Similarity scores between barcodes from CRISPR-edited embryos. **A**. Pairwise comparisons using cosine similarity of barcodes from untreated embryos (n = 14) as a control. **B-D**. Pairwise comparisons of barcodes from embryos targeted by single CRISPR systems; n = 8 each for LbaCas12a-edited only and SauCas9-edited only; n = 7 for SpyCas9-edited only. Based on the cosine similarity metric, LbaCas12a alone generates the most diverse barcodes. **E**. Pairwise comparisons of barcodes from triple-edited embryos (n = 35) fully saturating the orthogonal CRISPR system recorder.

**Figure S6.**
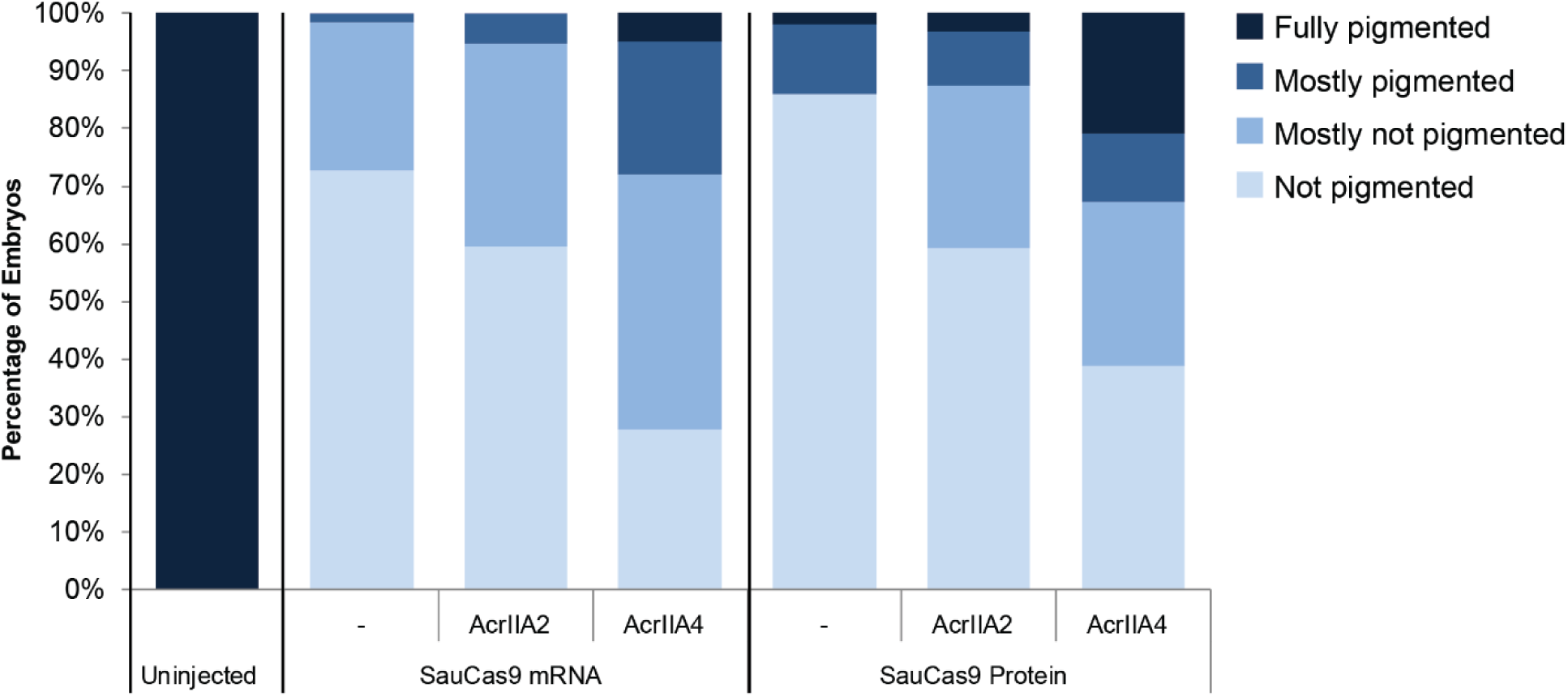
Anti-CRISPR inhibition of SauCas9 mRNA or protein microinjection. Percentage of embryos exhibiting pigmentation loss at 2 dpf after targeting the *tyr* gene with SauCas9, comparing SauCas9 mRNA or protein co-injection with anti-CRISPR mRNAs.

**Tables are in separate supplementary files**.

**Table S1: Addgene sources of CRISPR and anti-CRISPR plasmids used in this study**.

**Table S2: Oligos for cloning CRISPR and anti-CRISPR genes, and for in vitro transcription of sgRNAs**.

**Table S3: Raw data from microinjections**.

